# Where is the melody? Spontaneous attention orchestrates melody formation during polyphonic music listening

**DOI:** 10.1101/2025.08.26.672294

**Authors:** Martin M. Winchester, Kevin Reynolds, Charbel Nebo, Ian Cecil Scott, Giovanni M. Di Liberto

**Author notes:** Senior author (Giovanni M. Di Liberto).

## Abstract

Humans seamlessly process multi-voice music into a coherent perceptual whole. Yet the neural strategies supporting this experience remain unclear. One fundamental component of this process is the formation of melody, a core structural element of music. Previous work on monophonic listening has provided strong evidence for the neurophysiological basis of melody processing, for example indicating predictive processing as a foundational mechanism underlying melody encoding. However, considerable uncertainty remains about how melodies are formed during polyphonic music listening, as existing theories (e.g., divided attention, figure–ground model, stream integration) fail to unify the full range of empirical findings. Here, we combined behavioural measures with non-invasive electroencephalography (EEG) to probe spontaneous attentional bias and melodic expectation while participants listened to two-voice classical excerpts. Our uninstructed listening paradigm eliminated a major experimental constraint, creating a more ecologically valid setting. We found that attention bias was significantly influenced by both the high-voice superiority effect and intrinsic melodic statistics. We then employed transformer-based models to generate next-note expectation profiles and test competing theories of polyphonic perception. Drawing on our findings, we propose a weighted-integration framework in which attentional bias calibrates the overall degree of integration of the competing streams. In doing so, the proposed framework reconciles previous divergent accounts by showing that, even under free-listening conditions, melodies emerge through an attention-guided statistical integration mechanism.

**Highlights:**

1. EEG can be used to decode spontaneous attention during the uninstructed listening of polyphonic music.
2. Behavioural and neural data indicate that spontaneous attention is influenced by both high-voice superiority and melodic contour.
3. Attention bias impacts the neural encoding of the polyphonic streams, with strongest effects within 200 ms after note onset.
4. Stimuli that produced a stronger attention bias aligned with monophonic-model expectations, whereas stimuli with a weaker bias aligned with the Stream-Integration model.
5. We propose a bi-directional influence between attention and prediction mechanisms, with horizontal statistics impacting attention (i.e., salience), and attention impacting melody extraction.

## Introduction

Music from many cultures and traditions, including Western tonal music, typically leads to the perception of melody, which is a monophonic sequence of notes that a listener would sing, whistle, or hum back to identify the music (1, 2). While certain types of music are monophonic by nature (e.g., Sean-nós singing, nursery rhymes), leaving little doubt as to what the melody is, music typically involves multiple sound streams, or voices. A variety of factors, some acoustic (e.g., the particular instrument used, pitch distance) and others musical (e.g., meaningful melodic progressions), have been identified as key contributors to the perception of distinct sound streams in music. However, there remains considerable uncertainty on how those streams are processed by our brain into a coherent whole; and while melody has been proposed as a key structural element of music (3), the neural strategies underpinning its emergence from multi-stream compositions remains unclear.

While many theories have been proposed, one point of agreement is that attention mechanisms have a main role in polyphonic music perception. Much of the debate has centred around whether our brains are capable of dividing auditory attention across multiple auditory streams. In contrast with the widely studied multi-talker speech scenario, where attention can only be directed to one speaker or conversation at a time (4, 5), music is built so that distinct streams have strong relationships (e.g., harmonic) that can make them be perceived as coherent elements of a unified percept. Theories of polyphonic music perception have been formulated, for example explaining this phenomenon as the result of a divided attention between the streams (6, 7). A second view proposes that the attentional focus is directed to a single “foreground” stream, while also processing its harmonic relationship with the “background” (figure-ground model (8, 9)). As music unfolds, listeners may switch their attention to different streams, changing what is regarded as foreground and background. A third view proposes that our brains merge multiple music streams into a single complex melody, which then becomes the focus of attention (integration model (9)).

Tailored experiments have been used to test the validity of these models, often involving tasks such as error detection in which participants are asked to click a button when they hear anomalies in pitch, rhythm, or harmony (10). That work led to a wealth of knowledge on what our brains “can” do. However, these tailored tasks come at a cost, such as imposing tasks leading to neural processing strategies different from uninstructed listening. For example, error detection tasks may include the explicit instruction of focussing on one or all streams, leading to processing strategies (e.g., selective attention behaviour, rapid attention switching) that might or might not be employed during uninstructed listening. As such, while that work can certainly inform us on what our brains are capable of doing, it is less clear if that reflects the typical neural functioning during uninstructed listening.

Here, we investigate how the human brain processes two-stream polyphonic music during uninstructed listening by combining a behavioural task and non-invasive electroencephalography measurements (EEG). Participants were presented with classical music pieces synthesised with a high-quality virtual piano instrument, where each of the two voices consisted of a meaningful melody that would stand on its own (**Fig. 1**). In that condition, the listener’s attention was expected to primarily focus on the high pitch stream due to:

- the <u>high-voice superiority effect</u> (i.e., a bias towards the high-pitch stream; (11));
- and <u>the melody itself</u>, as the more attractive statistics are often placed at the higher pitch stream (11–14).

**Figure 1.**
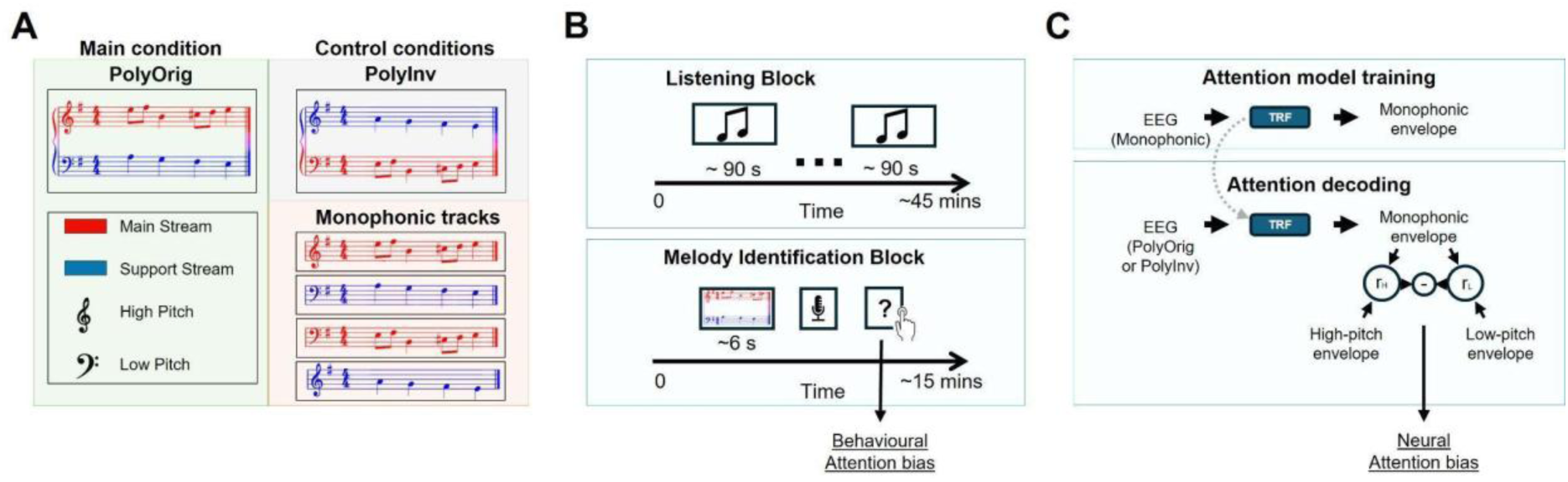
Experimental design. **(A)** Participants were presented with polyphonic music made of two monophonic streams (PolyOrig condition), with control stimuli where the average pitch heights for the two streams were inverted (PolyInv), and with the corresponding monophonic streams in isolation (Monophonic). All stimuli were synthesised from MIDI scores using high quality virtual piano sounds. **(B)** Each experimental session was organised into two blocks: a listening block, where EEG signals were recorded from participants during uninstructed music listening, followed by a music identification block. The music pieces were selected randomly from any condition, an a given piece was presented once, only in one of the three conditions, without repetitions. In the melody identification block, participants heard short polyphonic snippet from the same pieces, and were asked to sing back the melody and to indicate if that was the high or low pitch stream on a five-point scale. This led to a behavioural measurement of attention bias. **(C)** Attention bias was also measured from the neural signal. Attention decoding models were built on the monophonic condition, by fitting backward temporal response functions (TRF) on each participant to reconstruct the sound envelope from the EEG signal. TRF models were then applied to the polyphonic conditions to decode the attended melody, resulting in a neural measurement of attention bias.

To disentangle these two factors, alongside this main experimental condition involving the original polyphonic music (PolyOrig), our experiment included a control polyphonic condition where the average pitch-heights of the two-streams were inverted (PolyInv). An example of this manipulation is available at **Figure 1 Supplement 1**. Finally, we also included a monophonic condition including a random selection of the monophonic voices in both polyphonic conditions (Monophonic), which was used for training the models for the attention decoding analysis.

### Where is the attention?

First, we decode the attention bias in the listeners’ brains in the polyphonic listening conditions. In doing so, we answer the question: Where is the attended melody? To that end, we derived two attention bias scores. The first score is behavioural, and it reflects which of the two sound streams the listener sung back during a listen and repeat task in a dedicated block (behavioural attention bias score; **Figure 1**). The second score was derived from the EEG signal through an attention decoding procedure (neural attention bias score). Recent methodological developments provide us with tools to carry out that decoding from EEG signals based on regression methods (15). Specifically, we fit attention decoders on the EEG data recorded during the monophonic condition, which contained melodies selected from high and low pitch streams, and main and support streams. That way, the resulting decoders were unbiased with regard to pitch height and voice statistics, and were then applied to decode attention on the polyphonic conditions.

We hypothesised both music streams to be robustly encoded in the human cortex during polyphonic music listening, primarily reflecting the cortical encoding of low-level acoustic properties (e.g., sound envelope). With regard to auditory attention, the high-voice superiority effect and melody statistics are the two possible contributors to salience – timbre was not a factor in this experiment, as the sound stimuli were synthesised using MIDI sounds from a single instrument (see **Methods**). In the PolyOrig condition, both contributors were expected to primarily steer attention toward the high pitch stream by construction. Avoiding generalisations, the melodic material that characterises the main motif or primary thematic element, which is distinguished by its more intriguing and expressive features, is often constructed in the more exposed higher registers. For good practice, the main motif is supported by a harmonic and rhythmic structure constructed in order to enhance the attention on the main thematic material.

We designed the PolyInv condition to disentangle the two key contributors to saliency – high-voice superiority and melody statistics – by placing the main motif in the low-pitch voice. The pieces in this experiment, primarily double-counterpoint compositions, were selected specifically as the inversion of voices does not compromise their structural integrity. For example, this technique is a well-established practice in fugues. The bass (low-pitch) line, even if less ornamented than the main (high-pitch) melody, provides a fundamental harmonic structure. This harmonic foundation ensures that when the lines are inverted, the resulting music retains coherence and stability, even in the absence of a third voice to fully define the harmonic context. With this premise, three hypotheses were explored for the PolyInv condition: Hp0) High-voice superiority is the main contributor to attention bias, which would lead to a high-pitch bias; Hp1) The two contributors contrast each other, leading to small (or no) attentional bias; and Hp2) Melody statistics is the main contributor, meaning that PolyInv would lead to an inverted attention bias compared with PolyOrig, as the pitch lines for the two voices were swapped (**Fig. 2A**).

**Figure 2.**
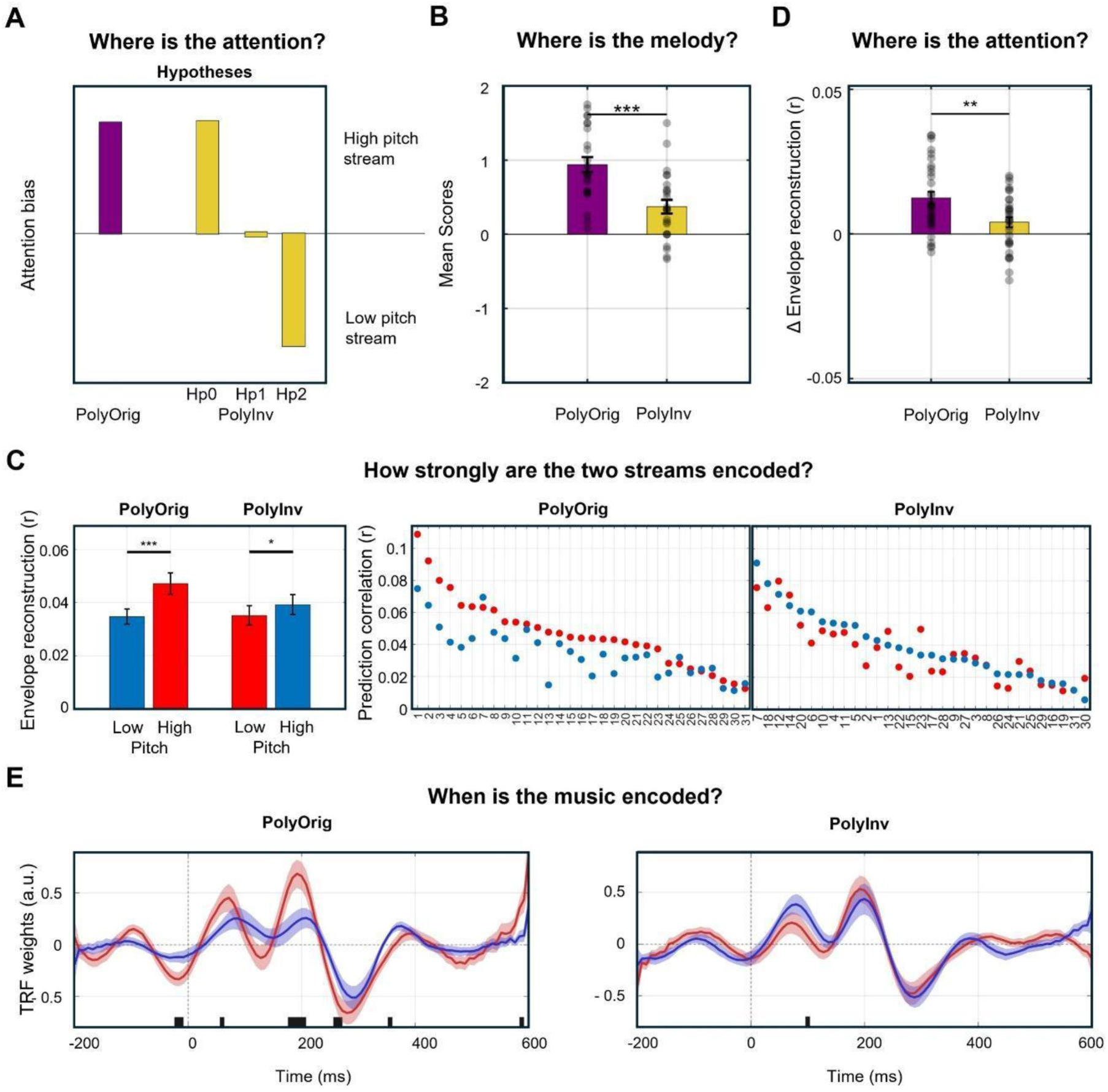
Behavioural and neural indices of attention bias during uninstructed listening of polyphonic music. **(A)** Illustration of the hypothesised attention bias scenarios for polyphonic music listening. A high-pitch bias was expected for PolyOrig by design. Dominant high-voice superiority effect or motif attractiveness were expected to lead to a strong attention bias toward the high (Hp0) and low (Hp2) pitch streams respectively, while a substantial reduction in attention bias would reflect a comparable contribution of the two factors (Hp1). **(B)** Behavioural result. The behavioural attention bias metric (mean ± SEM; ***p<0.001) was derived from the subjective reporting in the melody identification block. Subjective ratings indicated what stream was perceived as the main melody, from value -2 (low pitch) to value 2 (high pitch). **(C)** EEG decoding analysis. (Left) Envelope reconstruction correlations for the decoding analysis are reported (mean ± SEM; *p<0.05, ***p<0.001) for individual streams (high and low pitch) and conditions (PolyOrig and PolyInv). Colours refer to the motif (red: main melody; blue: support stream). Note that colours are inverted in the two conditions, reflecting the pitch inversion. (Right) Envelope reconstruction correlations for individual participants. **(D)** Neural attention bias index, which was obtained by subtracting the high and low pitch reconstruction correlations in (C) within each condition (Δcorrelation; mean ± SEM; **p<0.01). **(E)** Forward TRF model weights at channel Cz, providing insights into the temporal dynamics of the neural response to the two streams. Lines and shaded areas indicate the mean and SEM over participants respectively. Thick black lines indicate time points with statistically significantly difference across conditions.

### What is the melody?

After having established the attention bias of each participant and music piece, we present a second analysis to determine how the perceived melody is built by our brains. To that end, we rely on the known sensitivity of EEG signals to melodic expectations (10, 16–19). A music note may be more or less expected based on its prior context. Monophonic music listening leaves little doubt as to what that local context is, and models have been built that estimate next-note expectations, for example based on variable-order Markov Chains (20) and deep-learning architectures (21–23). Here, we use state-of-the-art transformer-based computational models of music to generate numerical hypotheses for our brains’ next-note expectations, matching different theories of polyphonic music perception. We then relate these simulations with neural signals recorded during the polyphonic conditions, determining which music processing strategy is most neurophysiologically plausible for uninstructed listening, and how attention bias relates with the selected strategy.

We considered a divided attention (6) and an integration model (9). The divided attention model would predict distinct melodic surprise for the two polyphonic streams, with equal strength. Measuring different strengths, in this two-stream listening scenario, would instead be compatible with a figure-ground model (7, 9), where the listening focus is directed to one particular stream (horizontal melody), which may alternate as the music unfolds, while the other is processed in function of the attended stream (vertical harmony). The other possibility that we considered was the integration model, according to which next-note expectations would be based on a single melody combining the two streams. Stream integration is in line with previous findings (9, 24) and compatible with the intuition that our vocal tract can only produce monophonic sequences. An auditory-motor neural pathway, condensing the auditory input into monophonic vocal motor commands, would be in line with the definition of melody itself (i.e., a monophonic sequence that a listener would hum back after hearing a music piece). Note that integration is not in contrast with the processing of other properties, such as vertical harmony, for example via a distinct neural pathway.

With that premise, our analysis aims to arbitrate among those three models. Measuring differences in the melody expectation encoding for the two streams in PolyOrig would contrast with the divided attention model. With regard to PolyInv, we had distinct hypotheses for the other two models. Specifically, the Figure-ground model PolyInv to impair the integration negatively, if anything, due to the alterations in vertical harmony produced by the pitch inversion. An integration model, instead, would be in line with an increased integration in PolyInv, where the attention bias was expected to be in-between the two streams (Hp1 in **Fig. 2A**).

## Results

Neural signals were recorded with EEG from 31 participants (16 males) during the uninstructed listening of polyphonic compositions or their monophonic components (*listening block;* **Fig. 1A**). The *listening block* involved the listening of monophonic and polyphonic pieces. In the second part of the experiment, participants undertook a melody identification task in a dedicated block, where short polyphonic music segments (∼4-8 seconds) were presented to the participants, who were asked to sing back the melody, and to indicate if they sung the high or low pitch stream on a five-point scale (see **Methods**). The analyses that follow integrate behavioural (**Fig. 1B**) and neural data (**Fig. 1C**) to address the following key questions: (a) Where was the focus of attention, and what factors influenced it? (b) What melody is encoded in the human brain during uninstructed listening to polyphonic music?

### Where is the attention? Investigating attention bias and its contributors using behavioural and neural measurements

In PolyOrig, the listeners’ attention was expected to focus on the high pitch stream by construction. PolyInv was built to disentangle the high-pitch superiority effect and the motif attractiveness. Measuring a strong attention bias toward the high-pitch stream in PolyInv would reflect a dominance of the high-pitch superiority effect (Hp0). Conversely, an attention bias toward the low-pitch stream in PolyInv would indicate a dominance of the motif attractiveness (Hp2). Our expectation was that both factors would contribute to the attention bias after the pitch inversion, thus leading to a reduced attention bias compared with PolyOrig (Hp1; **Fig. 2A**).

Behavioural measurements of attention bias confirm the high-pitch attention bias in the PolyOrig condition (*t*-test, *p* = 3.5*10^-9^, *d* = 1.88). The PolyInv condition also showed a high-pitch bias (*t*-test, *p* = 6.6*10^-4^, *d* = 0.80). Crucially, while the attention bias remained toward the high-pitch voice, its magnitude decreased with the voice inversion (paired *t*-test: *p* = 1.2*10^-4^, *d* = 0.95; **Fig. 2B**), in line with Hp1.

Next, we tested if the neural data also pointed to the same hypothesis. Neural measurements of attention bias were derived via an attention decoding analysis. Attention decoders were fit on the low-frequency EEG data (1-8 Hz) from the monophonic condition using lagged ridge regression, with a methodology referred to as backward temporal response function (TRF; (25–27)). The model fit identifies a linear combination of all EEG channels that produces an optimal reconstruction of the sound envelope. Since the monophonic listening condition only involved one music stream, participants could only attend to that stream, meaning that the resulting TRF model serves as an attention decoder. Sound envelope reconstructions were derived using that model on the EEG data from the polyphonic conditions. Pearson’s correlations were calculated between the reconstructed signal and the envelope of both polyphonic streams. The neural attention bias index was derived as the difference between those correlations (**Fig. 1C**).

Envelope reconstruction correlations showed a statistically significant attention bias (two-way ANOVA; main effect of stream: *F*(1,30) = 36.9, *p* = 1.1*10^-6^; **Fig. 2C**), no main effect of condition (PolyOrig vs. PolyInv: *F*(1,30) = 1.1, *p* = 0.296) and a statistically significant stream x condition interaction (*F*(1,30) = 8.8, *p* = 0.006; **Fig. 2D**). Post hoc tests confirmed, in line with the behavioural results and Hp1, a neural attention bias toward the high pitch stream in both PolyOrig (paired *t*-test: *p* = 2.4*10^-6^, *d* = 1.04) and PolyInv (*p* = 0.025, *d* = 0.42; **Fig. 2C**), and that the voice inversion led to a lower attention bias in PolyInv (paired *t*-test: *p* = 0.006, *d* = 0.53).

Further analyses were run to determine the impact of attention on the music stream encoding. Multivariate forward envelope TRFs were fit to build an optimal linear mapping from the envelope of the two input streams to the corresponding low-frequency EEG recording. Statistically significant differences were measured at the representative EEG channel Cz between the TRF weights for the two conditions (FDR corrected paired *t*-tests, *p*<0.05; **Fig. 2E**) in PolyOrig, with the effects emerging for all TRF components within the first 400ms after stimulus onset. Only a very short temporal cluster of significance emerged in PolyInv instead (∼80ms). Results were comparable at neighbouring channels such as Fz, while weaker effects emerged in more occipital scalp areas, such as Pz (not shown).

### What is the melody? Investigating melody formation by probing melodic prediction mechanisms in the listener’s brain

When asked to sing back a polyphonic piece, listeners produce a melody that is either one of the streams (segregation) or a mixture of the two (integration). The analysis that follows aimed at determining what the listeners’ brain regards as the melody. To that end, we relied on measurements of melodic expectations, which were previously validated in the context of monophonic music listening (16, 28, 29). Three models in the literature would predict different outcomes for this analysis: (a) the divided attention model predicts a simultaneous encoding of melodic expectations calculated on two streams as independent monophonic streams; (b) the figure-ground model would predict the encoding of melodic expectations for the main motif; and (c) the integration model would correspond to the encoding of melodic expectations for a melody that combines the two streams. Here, estimates of melodic expectations (surprise and entropy) for each melody stream were derived from an Anticipatory Music Transformer (23).

As a first step, we obtained a neural measurement of melodic expectation encoding on the monophonic condition, aiming to replicate previous findings in the literature (16, 30). The mapping between stimulus features and EEG is multivariate-to-multivariate, making methods such as the TRF (which is multivariate-to-univariate) suboptimal. Here we used Canonical Correlation Analysis (CCA) instead, because it accounts for all stimulus features and all EEG channels simultaneously. When comparing two expectation models, the difference in explained variance is typically very small, with the two models capturing essentially the same time-latencies and neural sources. As such, we opted for the method with the highest sensitivity in terms of explained variance, at the cost of model interpretability.

We measured match-vs-mismatch classification (**Fig. 3A**; (31, 32)), which informs on the EEG sensitivity to the music features, where higher classification scores indicate a stronger encoding of a given stimulus feature-set. Here, we compared the CCA mapping between two feature sets, one including only acoustic features (**A**: note onset, envelope, envelope derivative) and the other combining acoustic and melodic expectation features (**AM**, where M includes note onsets amplitude-modulated by pitch surprise and note onsets amplitude-modulated by pitch entropy for all streams in the stimulus). CCA models were fit for A and AM separately. As the match-vs-mismatch evaluation is a randomised procedure, we run 250 repetitions of that procedure to obtain a distribution of the participants average classification score for each model (A and AM). A statistically significant cortical encoding of melodic expectations was measured in the monophonic condition (AM > A, Wilcoxon rank sum: *p* < 10^-30^; **Fig. 3B-top**), consistent with previous EEG, MEG, and intracranial EEG work with Music Transformers and Markov models (16).

**Figure 3.**
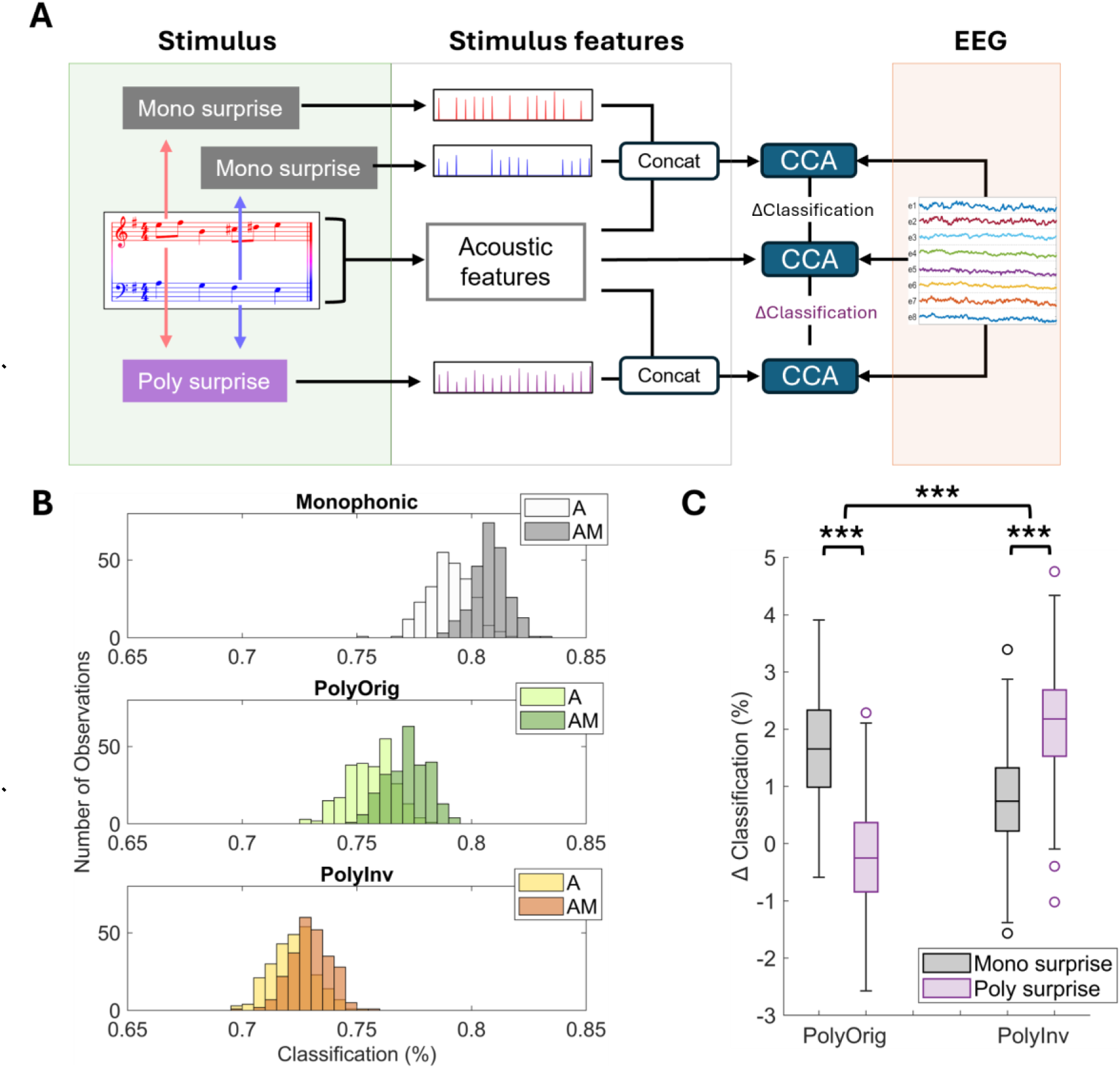
Low-frequency cortical encoding of melodic expectation during polyphonic music listening. **(A)** Schematics of the analysis method. Canonical Correlation Analysis (CCA) was run to study the stimulus-EEG relationship. Match-vs-mismatch classification scores were derived to quantify the strength of the neural encoding of a given stimulus feature-set. **(B)** The analysis was run separately for acoustic-only features (A) and acoustic+melodic expectations features (AM) in each of the three experimental conditions. The distributions in the figure indicate repetitions of the match-vs-mismatch procedure. **(C)** The gain in match-vs-mismatch classification after including the melodic expectation features, ΔClassification, is compared across conditions and models, informing on whether a monophonic or a polyphonic account of melodic expectations best fit the neural data. In the box-plot, the bottom and top edges mark the 25th and 75th percentiles respectively, while the mid-line indicates the median value (***p<0.001).

Next, we measured the cortical encoding of melodic expectation in the polyphonic conditions using the monophonic music transformer (**Fig. 3A, B**). A statistically significant encoding of melodic expectations emerged in both PolyOrig and PolyInv (two-way ANOVA; main effect of melodic expectations: *F*(1,249) = 425.3, *p* < 10^-30^; one-tailed *post hoc* Wilcoxon rank sum tests in PolyOrig and PolyInv: *p* < 10^-30^ and *p* = 1.6*10^-18^ respectively), with a significant main effect of condition (*F*(1,249) = 3967.5, *p* < 10^-30^). This analysis also detected a statistically significant condition x expectations interaction *F*(1,249) = 50.8, *p* = 1.1*10^-11^).

Finally, we tested if melodic expectations built with a polyphonic transformer better represented the EEG signal than for a monophonic model (**Fig. 3C**). A two-way ANOVA indicated main effects of model (monophonic vs. polyphonic: *F*(1,249) = 9.8, *p* = 0.002) and condition (PolyOrig vs. PolyInv: *F*(1,249) = 80.5, *p* = 7.2*10^-17^), as well as a statistically significant interaction effect (model x condition: *F*(1,249) = 357.6, *p* < 10^-30^). Interestingly, *post hoc* tests indicated that the EEG data more strongly related with the monophonic model in the PolyOrig condition and with the polyphonic model in the PolyInv condition (two-tailed *post hoc* Wilcoxon rank sum tests; PolyOrig:*p* = 2.7*10^-36^; PolyInv: *p* = 6.3*10^-26^).

While double-counterpoint music is particularly suitable for manipulations such as PolyInv, it is possible that the music transformer model’s prediction ability is affected nonetheless. As such, we carried out a tailored analysis to quantify the impact of that manipulation on the polyphonic model, finding a statistically significant increase in the surprise (KS test p = 9.7*10-130; D = 0.116), entropy (KS test p = 6.9*10-15; D = 0.0388), and perplexity (Wilcoxon signed-rank test p = 9.4*10-6, r = -0.783) for the polyphonic model. While our manipulation led to a reduced predictive ability of the model, it should be noted that these are very small effects (D<0.2).

The result in **Figure 3C** shows that the polyphonic model is a better predictor of EEG responses to PolyInv stimuli than the monophonic model, which is in line with our hypothesis. However, this effect may also be a reflection of a monophonic model that is affected by the pitch shift applied for the voice inversion, resulting in expectation values that are different from how the brain operates. We found that the pitch shift alters the calculation of the note surprise (Pearson’s correlation: *r* = 0.150, *p* = 7*10^-68^) and entropy (*r* = 0.106, *p* = 3*10^-34^). We re-run the CCA analysis on the PolyInv condition using expectation features for the monophonic model extracted from PolyOrig, thus excluding any influence of pitch shift. Interestingly, the key results in **Figure 3C** of a model x condition interaction also emerged in this control analysis (see **Fig. 3 Supplement 1**). Specifically, a two-way ANOVA indicated a main effects of model (monophonic vs. polyphonic: *F*(1,249) = 277.2, *p* = 2.5*10^-42^), no main effect of condition (PolyOrig vs. PolyInv: *F*(1,249) = 0.635, *p* = 0.426) and, crucially, a statistically significant interaction effect (model x condition: *F*(1,249) = 791.2, *p* = 2.9*\**10^-79^). *Post hoc* tests confirmed that the EEG data more strongly related with the polyphonic data model in the PolyInv condition (two-tailed *post hoc* Wilcoxon rank sum tests; PolyInv: *p* = 1.0*10^-85^).

## Discussion

This study investigated the neural processing of melody during polyphonic music listening. Previous research indicated that listeners can detect changes or errors in simultaneous music streams in some contexts (6, 8) but not others (9), challenging the possibility that our brains can truly divide attention between simultaneous melodic streams. Theories like the figure-ground model and stream integration (9) have been proposed as perceptual strategies that can compensate for the difficulty of truly dividing attention. While that work offers precious insights into what our brains “can” do, the tasks used previously for that research alter the listening experience, raising doubts with regard to how those models impact uninstructed listening. Here, we fill that gap by measuring the spontaneous attention bias in the listeners’ brains during the uninstructed listening of polyphonic music. First, we found that salience is affected by both the high-voice superiority effect and the statistical properties of the melodic lines (**Fig. 2**), providing a quantitative approach to disentangle the two effects. Second, we provide evidence for a weighted integration strategy, with the attention bias altering the stream integration rate (**Fig. 3**). Altogether, these results shed new light on the processing of melody during the uninstructed listening of polyphonic music, pointing to an inter-relationship between attention and statistical processing mechanisms, with attention orchestrating the way melody is formed.

### Investigating the contributors to attention during uninstructed polyphonic music listening

Attention is potentially the key mechanism for understanding how melody is processed during polyphonic music listening. To that end, we designed a paradigm where the attention focus would change across conditions, without the need for instructing the participants on where to direct their attentional focus. We opted for using double-counterpoint compositions involving two monophonic piano melodic lines, primarily from J.S. Bach (**Table 1**). As discussed by P.A. Scholes, such compositions involve melodic lines that “move in apparent independence and freedom though fitting together harmonically” (33), creating a scenario where both streams present attractive melodic progressions. As a counter example, let’s consider a scenario involving polyphonic music with two streams, where the support stream has infrequent notes with little melodic variation. That scenario would make the pitch-line inversion less effective, as the new high-pitch stream would not stand on its own as the main melody.

**Table 1:** Composers and titles of musical pieces used as stimuli.

| Composer | Title | Length | Composer | Title | Length |
| --- | --- | --- | --- | --- | --- |
| J. S. Bach | Bourrée in E minor | 1:47 | J. S. Bach | Invent 12 | 1:16 |
| J. S. Bach | Book 1 Fugue 02 | 1:57 | J. S. Bach | Invent 13 | 1:06 |
| J. S. Bach | Book 1 Fugue 10 | 1:31 | J. S. Bach | Invent 14 | 1:18 |
| J. S. Bach | Book 1 Prelude 02 | 1:51 | J. S. Bach | Invent 15 | 1:02 |
| J. S. Bach | Book 1 Prelude 03 | 1:21 | J. S. Bach | Invent 1 | 1:04 |
| J. S. Bach | Book 1 Prelude 14 | 1:22 | J. S. Bach | Invent 2 | 1:18 |
| J. S. Bach | Book 1 Prelude 24 | 1:57 | J. S. Bach | Invent 3 | 1:00 |
| J. S. Bach | Book 2 Fugue 02 | 2:08 | J. S. Bach | Invent 4 | 0:57 |
| J. S. Bach | Book 2 Fugue 07 | 2:35 | J. S. Bach | Invent 5 | 1:18 |
| J. S. Bach | Book 2 Prelude 02 | 2:18 | J. S. Bach | Invent 6 | 2:35 |
| J. S. Bach | Book 2 Prelude 07 | 1:38 | J. S. Bach | Invent 7 | 1:01 |
| J. S. Bach | Book 2 Prelude 12 | 2:06 | J. S. Bach | Invent 8 | 0:50 |
| C. P. E. Bach | La Caroline | 2:44 | J. S. Bach | Invent 9 | 1:08 |
| E. Grieg | Holberg Suite, Op. 40: III. Gavotte | 1:56 | I. Albeniz | España, Op.165, No. 2 | 2:16 |
| J. S. Bach | Invent 10 | 0:48 | J. Field | Nocturne No. 5 in B-flat Major | 1:27 |
| J. S. Bach | Invent 11 | 1:06 | J. S. Bach | Notebook2 16<br>March | 0:56 |

Previous studies demonstrated that the focus of selective auditory attention can be reliably decoded from EEG signals during sustained attention tasks (27) and, more recently, by studying instructed attention switches (34, 35). While that work was initially carried out on speech listening tasks in multi-talker scenarios (27, 36, 37), EEG attention decoding was also shown to be effective when considering polyphonic music during instructed selective attention tasks (15). Hausfeld and colleagues (38) went one step beyond by comparing an instructed selective attention task, where attention was steered to a given music stream, with a divided attention task, where participants were asked to focus on two music streams simultaneously. EEG attention decoding showed a preference toward the attended stream in the former, while no preference was measured in the latter task. Altogether, those results strongly support the reliability of decoding attention from EEG. However, that work focused on instructed listening, thus not informing us on how polyphonic music listening unfolds naturally. Other design choices further complicate the interpretation of those results, such as the use of different instruments for the distinct streams, which introduce another possible bias that, although interesting, was considered unnecessary to answer the research question in the present investigation. For that reason, our experiment only included virtual piano emulations generated from MIDI scores.

Our attention decoding analysis indicates that both high-pitch and melody statistics contribute to music salience. The present study teases apart two key factors influencing uninstructed attention by utilising two conditions involving the same type of music, with the sole difference that the pitch lines were inverted. Using behavioural (**Fig. 2B**) and EEG decoding indices (**Fig. 2D**), we found that the listeners’ steer their attention toward the high-pitch stream in both conditions, even though there was a substantially weaker bias in PolyInv. These results, together with the forward TRF results (**Fig. 2E**), are in line with our initial hypothesis that two key contributors are at play, and that the high-voice superiority effect is, in this case, stronger than the bias due to the melody attractiveness. It should be noted that our EEG results refer to the average attention bias throughout a piece, and that the behavioural index is only derived on a subsample of the music material. As such, that result would be compatible with different attention dynamics, such as a weaker attention bias throughout the piece, an increase in attention-switching between the voices, or their integration. Further to this, neural and behavioural measurement of attention bias were derived on different tasks, involving long (∼1.5 minutes) and short (∼4-8 seconds) music segments, with the latter happening as the second stage of the experiment, hence involving portions of stimuli that were heard in the first stage. As such, while we found consistent neural and behavioural attention bias measurements on average, further research should assess if differences in stimulus length, repetition, and task might affect the attention bias at more fine-grained levels (e.g., single participant and music piece).

Another limitation was the focus on a single music culture and style, which was a design choice allowing us to compare attention bias across two conditions that were minimally different. Indeed, different cultures are characterised by unique musical features. For example, some traditions employ distinct systems of note organisation, such as Makams, which incorporate additional pitches like quarter tones — which are not commonly found in Western music, except in certain avant-garde applications. Likewise, the inclusion of different musical styles would introduce varying melodic features, as they are constructed using diverse techniques. Future research should consider generalising the results of this research using a more diverse set of stimuli that can capture these interesting variabilities across music style and culture, as well as investigating the impact of other related factors, such as subjective biases (e.g., personal taste) and musical training.

This experiment involved two-stream stimuli where the main melody is primarily on the high pitch line, while both streams could stand on their own. The pitch inversion in PolyInv allowed us to study the neural encoding of music in conditions with different attention biases, while also disentangling the effects of high-pitch superiority and melody statistics. Switching pitch lines results in an inversion of the voicing, which affects the perception of the sound but not the harmonic context. Working with only two melodic streams facilitates their interchangeability, avoiding harmonic issues. Since a chord is typically defined by three notes, the absence of a third voice does not compromise harmonic clarity. In particular, many of these melodic elements, especially within the fugue, are constructed using the double-counterpoint technique, which guarantees the interchange of voices, without disrupting the structural integrity of the composition. Although the bass line is often less ornamented than the main melody, it provides essential harmonic grounding. This foundation ensures that even when the lines are inverted, the resulting music retains coherence and stability, despite the absence of a third voice to fully articulate the harmonic framework. As such, while generalisation the results of this study to other music styles requires further work, our choice of using double-counterpoint pieces enabled manipulations that were key for testing our hypotheses.

### Stream integration mechanisms contribute to melody formation

Numerous studies reported that neural responses to monophonic melodies reflect the expectation of a note, leading to patterns of neural activations that are compatible with a Bayesian view of the human brain, fitting frameworks such as predictive processing ((39); but see (21)). However, the relevance of those findings to the uninstructed listening of polyphonic music processing remains unclear. Listeners can identify a music piece by singing back its melody, indicating that part of polyphonic music processing involves transforming the incoming sound into a monophonic melody. This melody might correspond to a given stream or result from the integration of two or more streams. Previous work proposed theories on how that melody might be extracted. Here, we worked under the assumption that our brains process the resulting melody according to the predictive processing framework, where melodic expectations have been shown to be measurable with EEG (16). Using music transformers, we built numerical hypotheses on those melodic expectations, in one case assuming the processing of each monophonic stream (monophonic model), while the other model implements an integration of the two polyphonic streams (polyphonic model; **Fig. 3**).

Differently from existing theories, our results suggest that the human brain processes multiple music streams via a dynamic strategy. Specifically, a strong attention bias toward a dominant stream corresponded with a melody processing consistent with a monophonic processing of that stream. A weak attention bias, instead, corresponded with an integration of the two streams, as evidenced by a stronger alignment with the polyphonic model of melody expectations. In light of this result, we propose a *weighted-integration model*, in which attention modulates how strongly the two streams are integrated: with a strong bias, next-note prediction for the main stream are effectively monophonic (no integration); with a weaker bias, the support stream contributes proportionally (weighted integration). It should be noted that our results only speak to melody formation and melody prediction, and they do not address the important issue of how support streams are processed. While the figure-ground model offers a possible explanation, our data does not address that question nor exclude other possibilities, such as the divided attention model or one of its variations.

With the premise that attention might orchestrate music stream integration, one might observe that PolyOrig is more closely related with everyday listening of, for example, Western music, which typically involves a dominant melody. As a result, everyday music listening might be consistent with how the corresponding melodies would be processed in a monophonic context. This is a tantalising possibility, as it would make previous findings on monophonic melody perception directly relevant to more typical everyday multi-stream music listening. It should also be noted that our results involved averaging attention bias measurements throughout entire music pieces, while that bias likely changes within each piece. As such, the rate of integration might change dynamically at a more fine-grained level than when assessed here, calling for further investigation.

Our findings complement the past literature on simultaneous pitch sequences. Previous work using odd-ball paradigms measured mismatch-negativity responses to deviant notes in high or low pitch streams (40). That line of work, however, involves simplified paradigms including constraints making the task different from the typical music listening experience. We are especially referring to the use of deviant notes, very short stimuli, and isochronous note timing. Indeed, such previous work is valid and offers solid evidence of what our brains can do under those conditions. Our goal here was, instead, to determine how our brains operate in conditions one step closer to realistic music listening. As a result, we provide data confirming that melodic expectations can be measured in listening scenarios involving two-voice compositions, with results indicating that stream integration can naturally occur in correspondence of a weak attention bias.

## Conclusions

Additional research is necessary to determine if the inter-relationship between attention and melody formation is specific to this specific music style or if that is a general mechanism. Our methodology could be replicated on two-stream polyphonic music from other styles and cultures, with one caveat: the choice of an appropriate computational model of music must take into account the music culture or style of the stimuli and listener. That might constitute a limiting factor for under-represented music styles, even though the rapid developments in music transformer research are promising in that regard and may close that gap in the near future. Further work should also explore factors such as repetition, musical training, and music preference, which are expected to be particularly important for uninstructed listening tasks. Repetition, for example, may alter the attention bias and has been put forward as a possible way for disentangling the figure-ground and integration accounts of polyphonic music processing (9). With regard to musical training, while our results did not show any effects, our definition of musical training was quite varied in this sample. Future work aiming to shed light on that specific phenomenon should consider larger and more homogeneous samples.

In sum, we demonstrate that polyphonic music processing can be studied with EEG with an uninstructed listening task. Our findings include novel insights into the inter-relationship between attention and melody statistical processing. Based on these results, we propose an extension of the stream integration model of polyphonic music listening, where attention bias modulates the engagement of integration strategies. We also speculate that stream integration might be a consequence of an auditory-motor neural pathway condensing the auditory input into monophonic motor instructions that can be produced via our vocal tract. More work is necessary to validate and extend our findings and speculations, for example by considering more diverse sets of stimuli, music styles, and participant cohorts. Furthermore, while our study encapsulated melodic expectations into a single metric, our brains have been shown to encode expectation in relation to different attributes (16). And while this study only focussed on pitch and timing properties when studying expectations, it is possible that different results would emerge in relation with distinct properties. For example, it might be the case that expectations for low pitch stream melodies are more relevant to the rhythm, while pitch contour could be more important for high pitch streams, in line with previous results from basic auditory physiology research (41). Additional research with tailored tasks and stimuli should be explored to tackle those questions.

## Methods

### Participants

40 participants were recruited for this study. Data from nine participants were excluded due to technical issues with the synchronisation triggers in the EEG recording, leading to a dataset with 31 participants overall (16 males, aged between 19 and 30, mean = 23.2, std = 2.3), 8 of whom have had formal musical training or at least two years of professional musical experience. All participants gave their written informed consent to participate in this study. The study was undertaken in accordance with the Declaration of Helsinki and was approved by the ethics committee in the School of Psychology of Trinity College Dublin. Data collection was carried out between September 2022 and January 2024.

### Experimental design

Testing took place in a dark room. The experiment was organised into two experimental blocks: a listening block and a melody identification block (Figure 1), collecting EEG and behavioural measurements respectively. First, participants were presented to monophonic and polyphonic music pieces, with the sole instruction of listening to the music while looking at a fixation cross and minimising motor movement. After the listening block, the second part of the experiment involved a melody identification task, aided by a listen and repeat exercise. Specifically, participants were presented with short (4-8 seconds) segments of polyphonic music, extracted from previously presented PolyOrig or PolyInv pieces. After hearing a segment, participants were asked to sing it back. Then, they were presented with the two melodies as separate monophonic pieces with the order randomised and asked: “Which of the 2 melodies were you attempting to sing? 1: only the first melody; 2: mostly the first melody; 3: an equal mix of both; 4: mostly the second melody; 5: only the second melody”. EEG signals were not recorded during the second block and the recorded singing was not analysed. Importantly, participants sang back the melody before being presented with the monophonic streams, encouraging them to base their selection on their own melody extraction during polyphonic listening and reducing potential task-related influences, including presentation bias.

Neurobs Presentation software was used for coding the experiment (http://www.neurobs.com), which was carried out in a single session for all participants. Audio stimuli were presented at a sampling rate of 44,100 Hz and played through Sennheiser HD 280 Pro headphones. EEG data was simultaneously acquired with a sampling rate of 250 Hz, from twenty-four electrode positions using an mBrainTrain Smarting wireless system.

### Stimuli

Stimuli consisted of MIDI versions of 32 classical music pieces from a corpus of Bach and other Western composers (**Table 1**). The original pieces were ∼150s long snippets of polyphonic music. Minor manual corrections were applied to ensure that each piece was the combination of two monophonic streams, ensuring no more than two notes co-occurred at any given time. These two streams were then separated based into the high-pitch stream and the low-pitch stream (corresponding to the main and support stream respectively). These original polyphonic pieces (PolyOrig) were then manipulated to generate stimuli for other two conditions: PolyInv and monophonic. The two streams were pitch-shifted (typically by one octave) to invert the pitch lines, ensuring that all notes in the low-pitched stream were below the high-pitched stream. This ensured that the high- and low-pitch streams were clearly distinguishable in terms of pitch range (42). This manipulation led to the PolyInv stimuli, where the melody originally built to be the main stream was now at the low pitch line. Finally, the monophonic stimuli consisted of the individual streams extracted from PolyOrig and PolyInv. Finally, the music integrity of the resulting pieces was verified by a music expert and composer (I.C.S.).

### EEG data preprocessing

EEG and stimulus data were anonymised after collection and were stored using the Continuous-event Neural Data (CND) format. EEG data was preprocessed and analysed offline using MATLAB R2023a software using custom code built starting from the analysis scripts of the CNSP open science initiative (*github.com/CNSP-Workshop/CNSP-resources*). EEG signals were filtered using high and low pass filters at cut off frequencies 1 and 8 Hz respectively. All filtering was done with Butterworth zero-phase filters of order two, implemented with the *filtfilt* function. EEG signals were downsampled to 125 Hz. EEG channels with a variance exceeding three times that of the surrounding ones were replaced by an estimate calculated using spherical spline interpolation. EEG data were re-referenced to the global average of all channels.

### Temporal Response Function

A system identification approach was used to compute the channel-specific mapping between music features and EEG responses. This method, referred to as the temporal response function (TRF), models the neural response at time *t* and channel *η* as a linear convolution of the stimulus property *s*(*t*) with an unknown, channel-specific filter *w*(*τ*, *η*), plus a residual term:

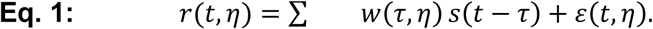

Here, *r*(*t*,η) is the instantaneous neural response, and ε(*t*, *η*) is the residual error not explained by the model.

The TRF *w*(*τ*,*η*) was estimated using regularised linear regression (ridge regression) to minimise overfitting. The solution is given by:

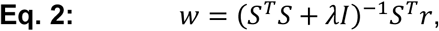

where *S* is the lagged time series matrix of the stimulus features, *λ* is the ridge parameter, and *I* is the identity matrix.

Model performance is evaluated using leave-one-out cross-validation across trials. The quality of a prediction is quantified by calculating Pearson’s correlation between the pre-processed recorded signals and the corresponding predictions at each scalp electrode.

Conversely, the backward TRF is a linear filter that can be fit to decode stimulus features from the neural recording. To estimate spontaneous attention, a backward TRF model is fit on the monophonic condition, where only one stream is played at a time. Note that the monophonic stimuli in the experiment were randomly selected from all streams in any of the polyphonic conditions, constructing a balanced training set and leading to a decoder that reconstructs the envelope of the attended melody. That decoder was then applied to the polyphonic conditions, producing envelope reconstructions that best correlate with the attended stream, which is then used to compare the decoding accuracies of the two streams the participants actually heard.

### Canonical Correlation Analysis

The comparison between the polyphonic expectation models was carried out using Canonical Correlation Analysis (CCA), which implements a multivariate-to-multivariate mapping between the stimulus features and the EEG signal. The main advantage over TRFs is that all stimulus features and EEG dimensions are related by applying linear transformations to both stimulus and EEG dimensions, projecting them onto a shared canonical subspace, leading to higher correlation values.

In its more general definition, given two sets of multichannel data X and Y of size T × J_1_ and T × J_2_, CCA finds linear transformations of both that make them maximally correlated. Specifically, CCA produces the transformation matrices W_1_ and W_2_ (sizes J_1_ × J_0_ and J_2_ × J_0_, where J_0_ < min(J_1_,J_2_)) that maximise the correlation between pairs of columns of XW_1_ and YW_2_, while making the columns of each transformed data matrix mutually uncorrelated. The first pair of canonical components (CC) is the linear combination of X and Y with highest possible correlation. The next pair of CCs are the most highly correlated combinations orthogonal to the first, and so-on. When X and Y represent the stimulus features and neural data respectively, CCA captures their *instantaneous* relationship. As such, similar to the TRF analysis, CCA can be used to study time-lagged relationships i.e., where the neural reaction to a given sound input unfolds with a non-zero delay and duration. Previous research determined that this modelling approach is effective at measuring sound-EEG mappings, providing different ways to include time-lags in the CCA modelling. Here we have done so by including time-lagged versions of both stimulus and EEG in X and Y respectively (please refer to the CCA-2 implementation in (36)), including a lag window of 0 to 500 ms. This leads to a substantial expansion of the X and Y matrices. To prevent overfitting, both X and Y were reduced to 24 dimensions via principal component analyses. The CCA results were evaluated with leave-one-out cross validation to control for overfitting, exploring input-output shifts over a window of -500 to 500ms.

The neural encoding of a given set of features was evaluated with the match-vs-mismatch accuracy metric. Stimulus and EEG time-series are partitioned into 20-second segments. Each segment is projected onto the canonical space. Correlations are calculated between the stimulus features and two EEG segments in the canonical space: the corresponding EEG segment (match) and a randomly selected one (mismatch). A binary classification accuracy score is derived by counting the proportion of match > mismatch correlations across all segments (31), providing a measurement of stimulus feature sensitivity for the EEG signal of a given participant. This metric is adopted in the analysis in Figure 3 to determine if monophonic or polyphonic expectation features match neural activity more closely in a given experimental condition.

### Anticipatory Music Transformer

AMT is a pretrained causal transformer model for symbolic music. Transformer architectures internally build probability distribution functions for each upcoming note based on long term memory (training set) and short term memory (context). We used the stanford-crfm/music-large-800k checkpoint, which was pretrained on large-scale music corpora including Lakh MIDI, MetaMidi, FMA transcriptions, and approximately 450,000 commercial recordings transcribed automatically. AMT represents symbolic music as sequentially ordered tokens encoding pitch and onset of each note individually and was trained to predict each upcoming token given the preceding context. Thus notes that occur simultaneously are encoded as separate MIDI events that share the same onset. The released model uses a context length of 1024 tokens.

### Stimuli feature extraction

Acoustic and melodic features were extracted from the audio files. Acoustic features included the sound envelope (*Env*), which was extracted from the Hilbert transform of the acoustic waveform. Second, we included the halfway rectified envelope derivative (*Env’*), which was calculated on the Hilbert envelope at the original sampling rate (44,100 Hz), before downsampling both *Env* and *Env’*. Melodic features were derived using AMT. The melodic features used were the surprise and entropy of the pitch of the notes (*S*, *H*), extracted from the model’s conditional probability mass function over the set of possible notes. Surprise is the information content of the note.

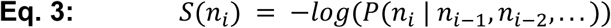

Where *n_i_* is the current note and *n_i-1_, n_i-2_*, … are the previous notes. The entropy represents the uncertainty at the time of the new note, calculated as the Shannon entropy over the distribution of the set of notes (*N*).

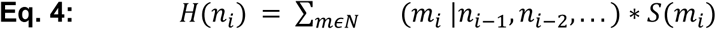

For the polyphonic model, which may generate expectation values for multiple simultaneous notes, we utilised the last expectation value at a given timepoint. This ensures the event is calculated within the context of the simultaneous note, whereas the first event would appear as if a single note were being played.

All features were downsampled to 125 Hz for analysis.

### Statistical analysis

All statistical analyses were carried out in MATLAB R2023a. Analyses directly comparing the groups were performed using repeated measures two-way ANOVAs with the *ranova* function. One-sample *t*-tests were used for post hoc tests. Correction for multiple comparisons was applied where necessary via Benjamini-Hochberg Procedure with the *fdr_bh* function. Effect sizes are reported using Cohen’s *d*.

## Supporting information

Supplemental wav

## Data availability

Analysis code and data (both EEG signals and stimuli) are publicly available in a standardised format (Continuous-event Neural Data (43)) on OSF at this link: https://osf.io/bjdh6

## Author contributions

Conceptualisation, funding acquisition, supervision, and project administration: GDL. Experimental design, pilot data collection and preliminary analyses: KR, CN, and GDL. Data collection and data curation: MW. Methodology, formal data analysis, and visualisation: MW and GDL. Supervision and project administration: GDL. Validation of music stimuli and musicology implications: ICS. Writing of first draft: GDL and MW. Editing & Review: All authors.

## Acknowledgments

This research was conducted with the financial support of Research Ireland at ADAPT, the Research Ireland Centre for AI-Driven Digital Content Technology at Trinity College Dublin and University College Dublin [13/RC/2106_P2]. I.C.S. was supported by a Government of Ireland Postgraduate Scholarship (Irish Research Council). For the purpose of Open Access, the authors have applied a CC BY public copyright licence to any Author Accepted Manuscript version arising from this submission. We thank the Cognition and Natural Sensory Processing (CNSP) initiative, which provided the blueprint for the analysis code and data standardisation guidelines used in this work. We thank Asena Akkaya and Amirhossein Chalehchaleh for their help with data collection.

---

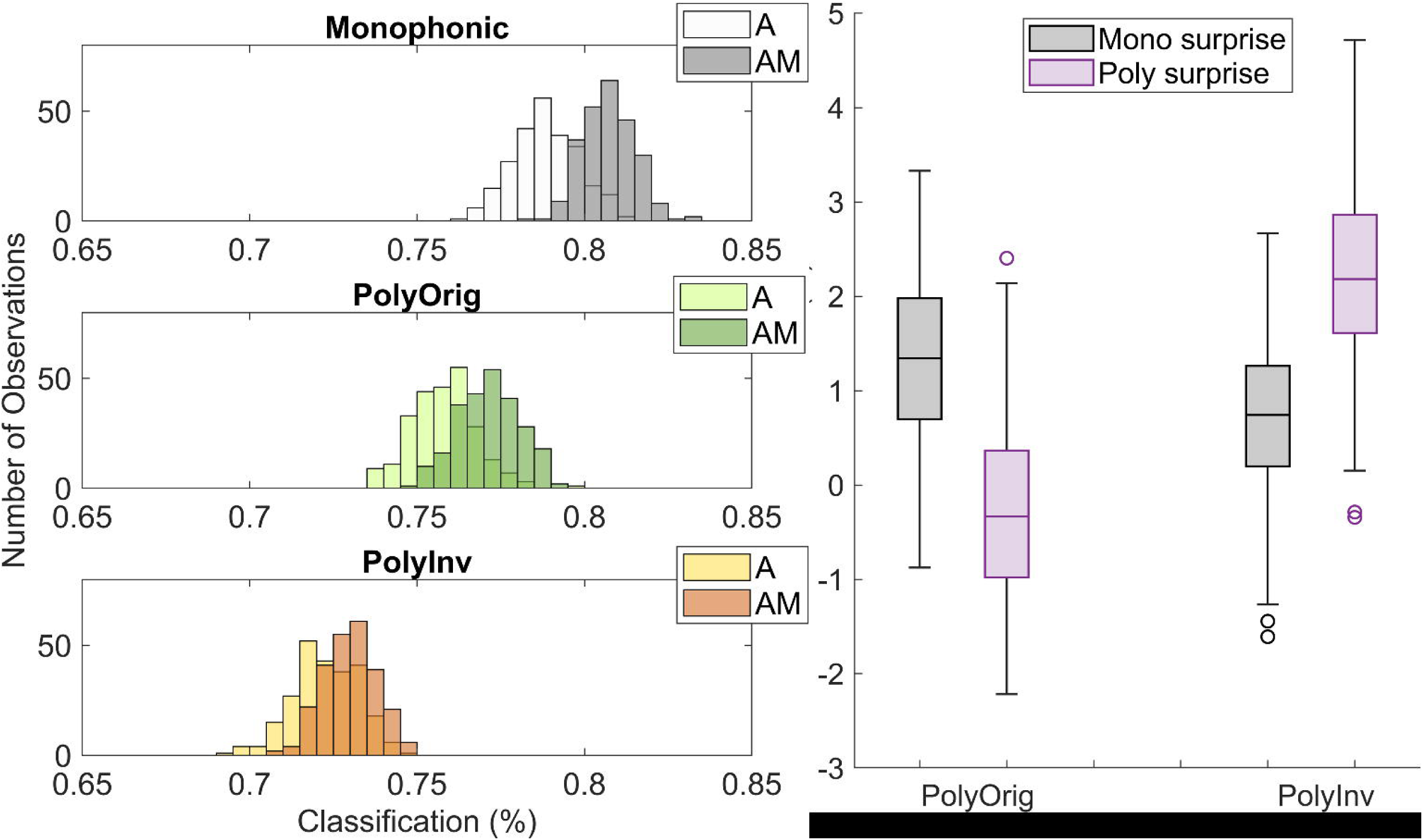

## References

1. Salamon J, Gómez E. Melody extraction from polyphonic music signals using pitch contour characteristics. IEEE transactions on audio, speech, and language processing. 2012;20(6):1759–70.

2. Poliner GE, Ellis DP, Ehmann AF, Gómez E, Streich S, Ong B. Melody transcription from music audio: Approaches and evaluation. IEEE Transactions on Audio, Speech, and Language Processing. 2007;15(4):1247–56.

3. Dowling W, Harwood D. Music Cognition Academic Press. London; 1986.

4. Raghavan VS, O’Sullivan J, Bickel S, Mehta AD, Mesgarani N. Distinct neural encoding of glimpsed and masked speech in multitalker situations. Plos Biology. 2023;21(6):e3002128.

5. Park H, Gross J. Get the gist of the story: Neural map of topic keywords in multi-speaker environment. bioRxiv. 2022:2022.05. 05.490770.

6. Gregory AH. Listening to polyphonic music. Psychology of Music. 1990;18(2):163–70.

7. Dowling WJ. The perception of interleaved melodies. Cognitive Psychology. 1973;5(3):322–37.

8. Sloboda J, Edworthy J. Attending to two melodies at once: the of key relatedness. Psychology of Music. 1981;9(1):39–43.

9. Bigand E, McAdams S, Forêt S. Divided attention in music. International Journal of Psychology. 2000;35(6):270–8.

10. Crawley EJ, Acker-Mills BE, Pastore RE, Weil S. Change detection in multi-voice music: the role of musical structure, musical training, and task demands. Journal of Experimental Psychology: Human Perception and Performance. 2002;28(2):367.

11. Trainor LJ, Marie C, Bruce IC, Bidelman GM. Explaining the high voice superiority effect in polyphonic music: Evidence from cortical evoked potentials and peripheral auditory models. Hearing Research. 2014;308:60–70.

12. Marie C, Fujioka T, Herrington L, Trainor LJ. The high-voice superiority effect in polyphonic music is influenced by experience: A comparison of musicians who play soprano-range compared with bass-range instruments. Psychomusicology: Music, Mind, and Brain. 2012;22(2):97.

13. Marie C, Trainor LJ. Early development of polyphonic sound encoding and the high voice superiority effect. Neuropsychologia. 2014;57:50–8.

14. Marie C, Trainor LJ. Development of simultaneous pitch encoding: infants show a high voice superiority effect. Cerebral Cortex. 2013;23(3):660–9.

15. Cantisani G, Essid S, Richard G, editors. EEG-based decoding of auditory attention to a target instrument in polyphonic music. 2019 IEEE workshop on applications of signal processing to audio and acoustics (WASPAA); 2019: IEEE.

16. Di Liberto GM, Pelofi C, Bianco R, Patel P, Mehta AD, Herrero JL, et al. Cortical encoding of melodic expectations in human temporal cortex. Elife. 2020;9:e51784.

17. Pearce MT. Statistical learning and probabilistic prediction in music cognition: mechanisms of stylistic enculturation. Annals of the New York Academy of Sciences. 2018;1423(1):378–95.

18. Bianco R, Harrison PMC, Hu M, Bolger C, Picken S, Pearce MT, et al. Long-term implicit memory for sequential auditory patterns in humans. eLife. 2020;9:e56073.

19. Trainor LJ, Zatorre RJ. The neurobiological basis of musical expectations. The Oxford handbook of music psychology. 2009:171–83.

20. Pearce MT, Wiggins GA. Auditory Expectation: The Information Dynamics of Music Perception and Cognition. Topics in Cognitive Science. 2012;4(4):625–52.

21. Robert P, Van Cang MP, Mercier M, Trébuchon A, Bartolomei F, Arnal LH, et al. Multi-stream predictions in human auditory cortex during natural music listening. bioRxiv. 2024:2024.11.27.625704.

22. Oore S, Simon I, Dieleman S, Eck D, Simonyan K. This time with feeling: Learning expressive musical performance. Neural Computing and Applications. 2020;32:955–67.

23. Thickstun J, Hall D, Donahue C, Liang P. Anticipatory music transformer. arXiv preprint arXiv:230608620. 2023.

24. Disbergen NR, Valente G, Formisano E, Zatorre RJ. Assessing top-down and bottom-up contributions to auditory stream segregation and integration with polyphonic music. Frontiers in neuroscience. 2018;12:121.

25. Crosse MJ, Di Liberto GM, Bednar A, Lalor EC. The multivariate temporal response function (mTRF) toolbox: A MATLAB toolbox for relating neural signals to continuous stimuli. Frontiers in Human Neuroscience. 2016;10(NOV2016).

26. Crosse MJ, Zuk NJ, Di Liberto GM, Nidiffer AR, Molholm S, Lalor EC. Linear Modeling of Neurophysiological Responses to Speech and Other Continuous Stimuli: Methodological Considerations for Applied Research. Frontiers in neuroscience. 2021;15:705621-.

27. O’Sullivan JA, Power AJ, Mesgarani N, Rajaram S, Foxe JJ, Shinn-Cunningham BG, et al. Attentional Selection in a Cocktail Party Environment Can Be Decoded from Single-Trial EEG. Cerebral Cortex. 2014:bht355-bht.

28. Di Liberto GM, Marion G, Shamma SA. The Music of Silence: Part II: Music Listening Induces Imagery Responses. The Journal of Neuroscience. 2021;41(35):7449.

29. Marion G, Di Liberto GM, Shamma SA. The Music of Silence. Part I: Responses to Musical Imagery Accurately Encode Melodic Expectations and Acoustics. Journal of Neuroscience. 2021.

30. Kern P, Heilbron M, de Lange FP, Spaak E. Cortical activity during naturalistic music listening reflects short-range predictions based on long-term experience. eLife. 2022;11:e80935.

31. de Cheveigné A, Di Liberto GM, Arzounian D, Wong DDE, Hjortkjær J, Fuglsang S, et al. Multiway canonical correlation analysis of brain data. NeuroImage. 2019;186:728–40.

32. Di Liberto GM, Wong D, Melnik GA, de Cheveigne A. Low-frequency cortical responses to natural speech reflect probabilistic phonotactics. NeuroImage. 2019;196:237–47.

33. Scholes PA. In true polyphonic music every line develops its own metric pattern without even a regularity of accented beats among voices. The uniformity is brought about in the harmony alone.: London: Oxford Univ. Press; 1938. 742 p.

34. Hjortkjær J, Wong DD, Catania A, Märcher-Rørsted J, Ceolini E, Fuglsang SA, et al. Real-time control of a hearing instrument with EEG-based attention decoding. Journal of Neural Engineering. 2025;22(1):016027.

35. Haro S, Rao HM, Quatieri TF, Smalt CJ. EEG alpha and pupil diameter reflect endogenous auditory attention switching and listening effort. Eur J Neurosci. 2022;55(5):1262–77.

36. De Cheveigné A, Wong DDE, Di Liberto GM, Hjortkjær J, Slaney M, Lalor E. Decoding the auditory brain with canonical component analysis. NeuroImage. 2018;172:206–16.

37. Fuglsang SA, Dau T, Hjortkjær J. Noise-robust cortical tracking of attended speech in real-world acoustic scenes. NeuroImage. 2017;156:435–44.

38. Hausfeld L, Disbergen NR, Valente G, Zatorre RJ, Formisano E. Modulating cortical instrument representations during auditory stream segregation and integration with polyphonic music. Frontiers in neuroscience. 2021;15:635937.

39. Clark A. Surfing Uncertainty: Prediction, Action, and the Embodied Mind: Oxford University Press; 2016.

40. Fujioka T, Trainor LJ, Ross B, Kakigi R, Pantev C. Automatic encoding of polyphonic melodies in musicians and nonmusicians. Journal of cognitive neuroscience. 2005;17(10):1578–92.

41. Hove MJ, Marie C, Bruce IC, Trainor LJ. Superior time perception for lower musical pitch explains why bass-ranged instruments lay down musical rhythms. Proceedings of the National Academy of Sciences. 2014;111(28):10383–8.

42. Huron D. The Avoidance of Part-Crossing in Polyphonic Music: Perceptual Evidence and Musical Practice. Music Perception: An Interdisciplinary Journal. 1991;9(1):93–103.

43. Di Liberto GM, Nidiffer A, Crosse MJ, Zuk N, Haro S, Cantisani G, et al. A standardised open science framework for sharing and re-analysing neural data acquired to continuous stimuli. Neurons, Behavior, Data analysis, and Theory. 2024:1–25.

